## Supplementary material for "Socially regulated developmental plasticity in the color pattern of an anemonefish": Figure S1

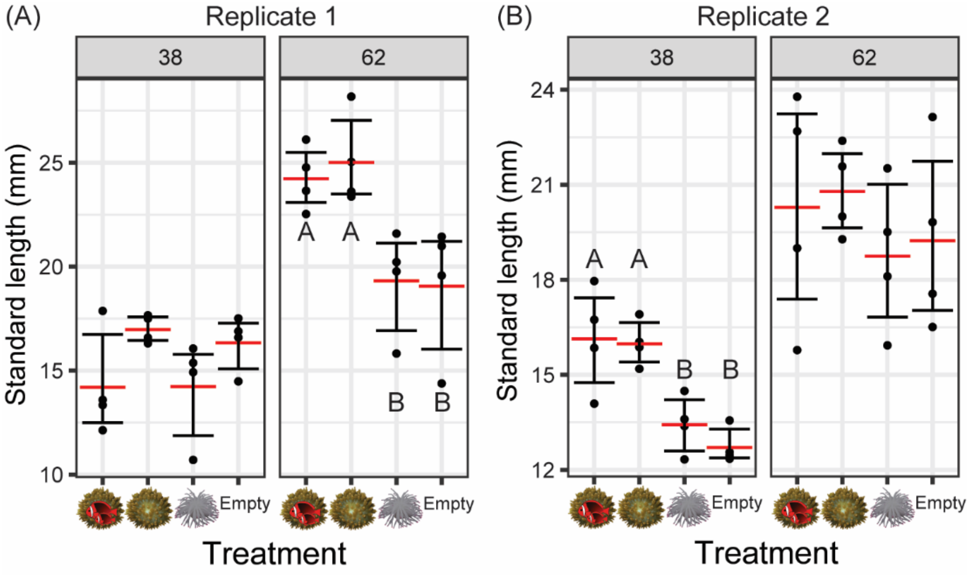


**Supplementary Figure 1. (A)** Juvenile standard length (mm) per environmental treatment at 38 and 62 days-post-hatch/dph in replicate 1 and **(B)** replicate 2. Letters denote statistical significance (ANOVA, $p_{adj} < 0.05$) grouping.
