## Supplementary material for "Socially regulated developmental plasticity in the color pattern of an anemonefish": Figure S2

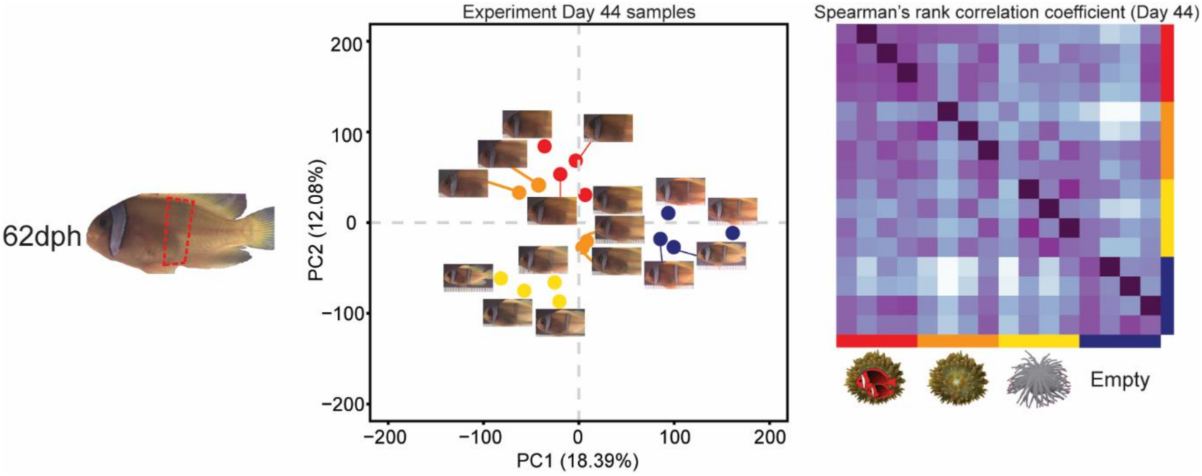


**Supplementary Figure 2.** PCA showing the distribution of Day 44 (62dph) samples along PC1 and PC2 according to total gene expression, along with correlation matrix depicting pairwise Spearman's rank correlation coefficients, calculated using transcriptomic data.
