## Supplementary material for "Socially regulated developmental plasticity in the color pattern of an anemonefish": Figure S3

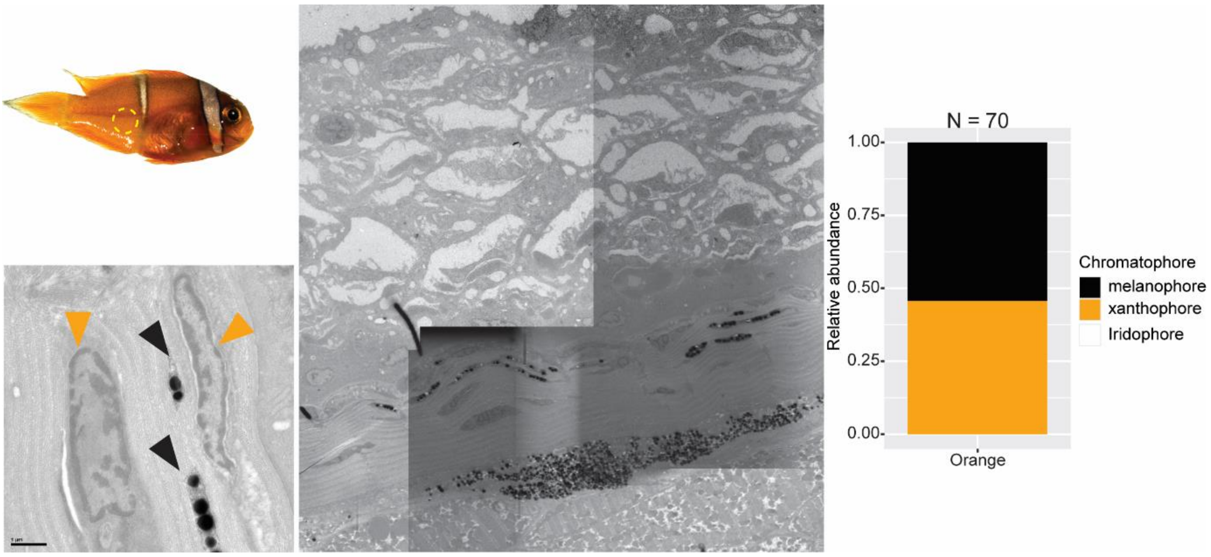


**Supplementary Figure 3.** Example of a TEM section in the orange skin peripheral to the fading white bar, with a closeup showing pigment cells (colored triangles). Bar plot shows the proportional abundance (counts) of different chromatophore types counted from 70 cells in three sections.
