## Supplementary material for "Socially regulated developmental plasticity in the color pattern of an anemonefish": Figure S4

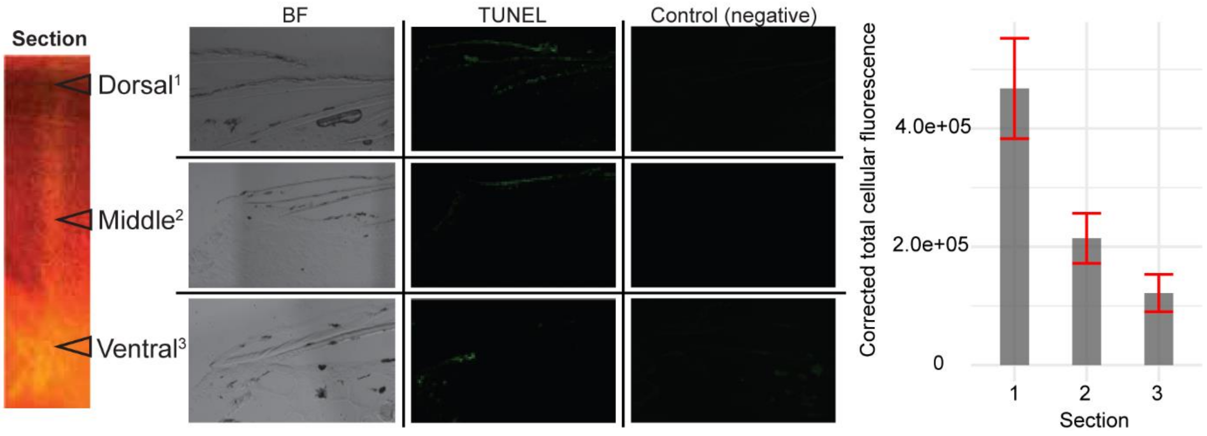


**Supplementary Figure 4.** TUNEL assay images taken in three sectioned (10µm thick) dorsoventral regions of the fading body bar of *A. frenatus*, including brightfield 'BF', TUNEL reaction mix treated, and negative control. The side bar plot depicts the mean (±0.95 CI) area corrected total cellular fluorescence (CTCF) measured from individual cells across three sections per the ventral (section 1, n = 64), middle (section 2, n = 75), and dorsal (section 3, n = 63) regions.
