## Supplementary material for "Socially regulated developmental plasticity in the color pattern of an anemonefish": Figure S5

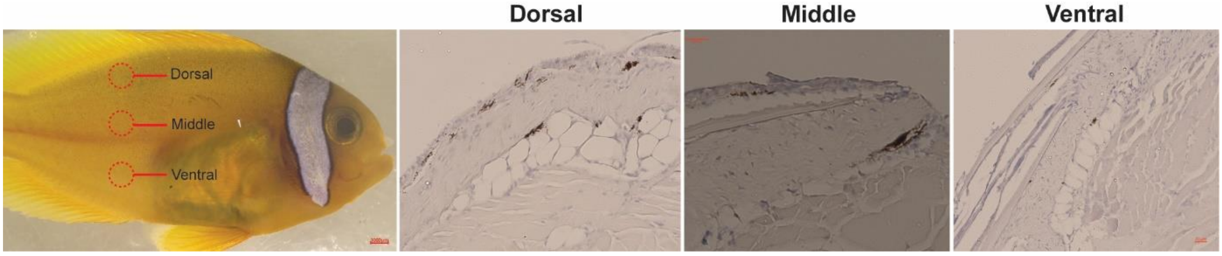


**Supplementary Figure 5.** Micrographs of immunostained sections using cleaved/activated Caspase-3 antibody with DAB in the late stage, faded white skin. No DAB+ cells were observed.
