## Supplementary material for "Socially regulated developmental plasticity in the color pattern of an anemonefish": Figure S6

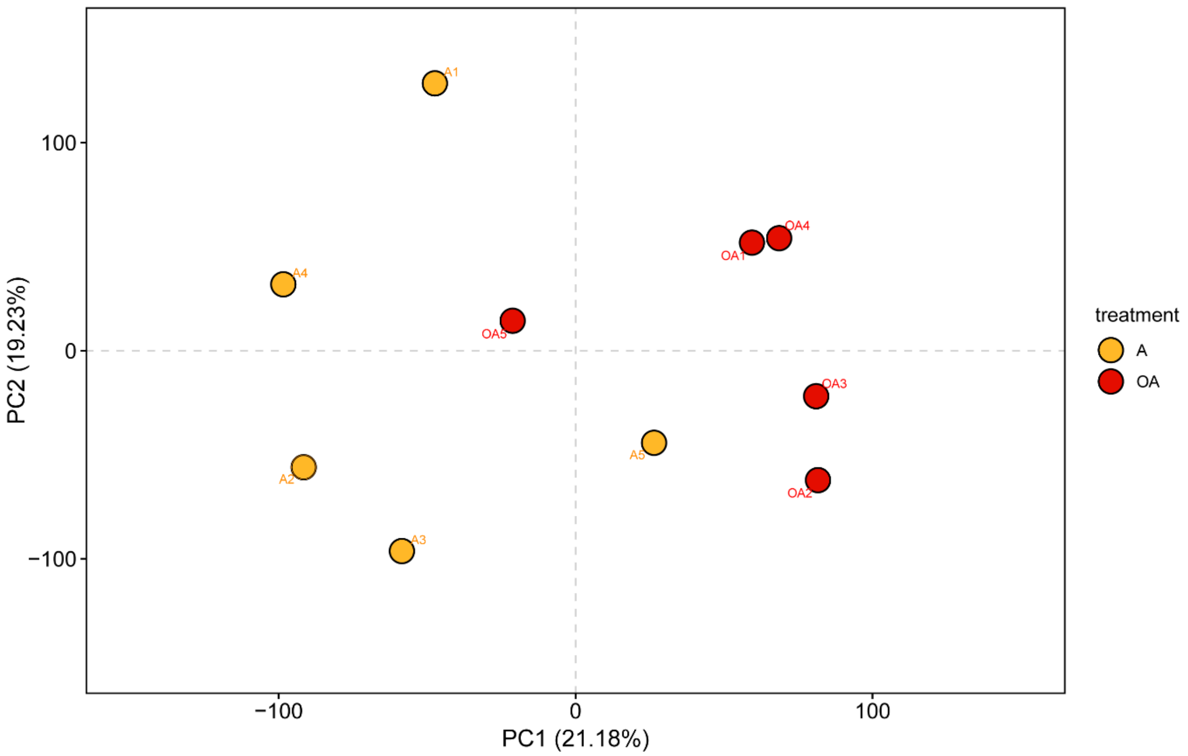


**Supplementary Figure 6.** PCA showing distribution of whole-body total transcriptomes for juvenile *A. frenatus* (n = 5) which cohabitated with or without an adult pair.
