## Supplementary material for "Socially regulated developmental plasticity in the color pattern of an anemonefish": Figure S7

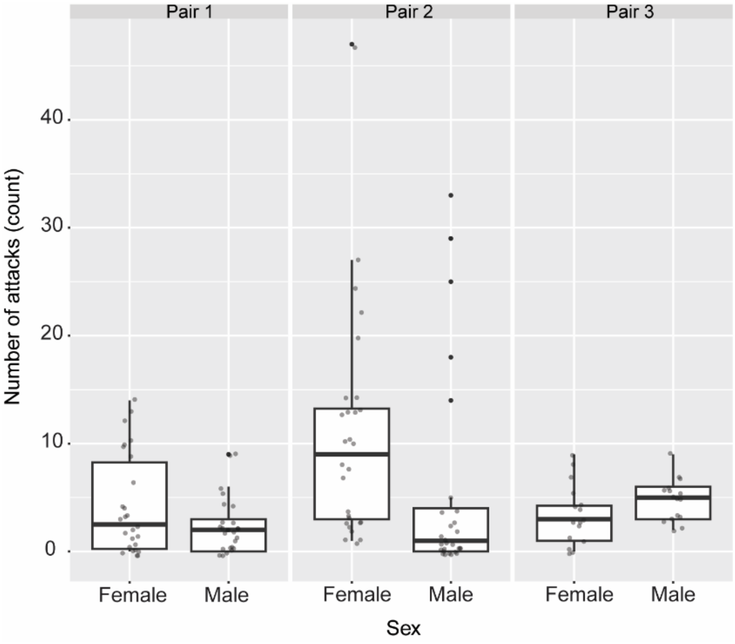


**Supplementary Figure 7.** Individual counts of attacks made by male and female adults per *A. frenatus* pair presented with juvenile anemonefish. Presented is the combined behavior data from Experiments 1 and 2. Boxes represent the median, 25th and 75th percentiles, and range (whiskers). Points represent the summed number of attacks per trial.
