## Supplementary material for "Socially regulated developmental plasticity in the color pattern of an anemonefish": Figure S8

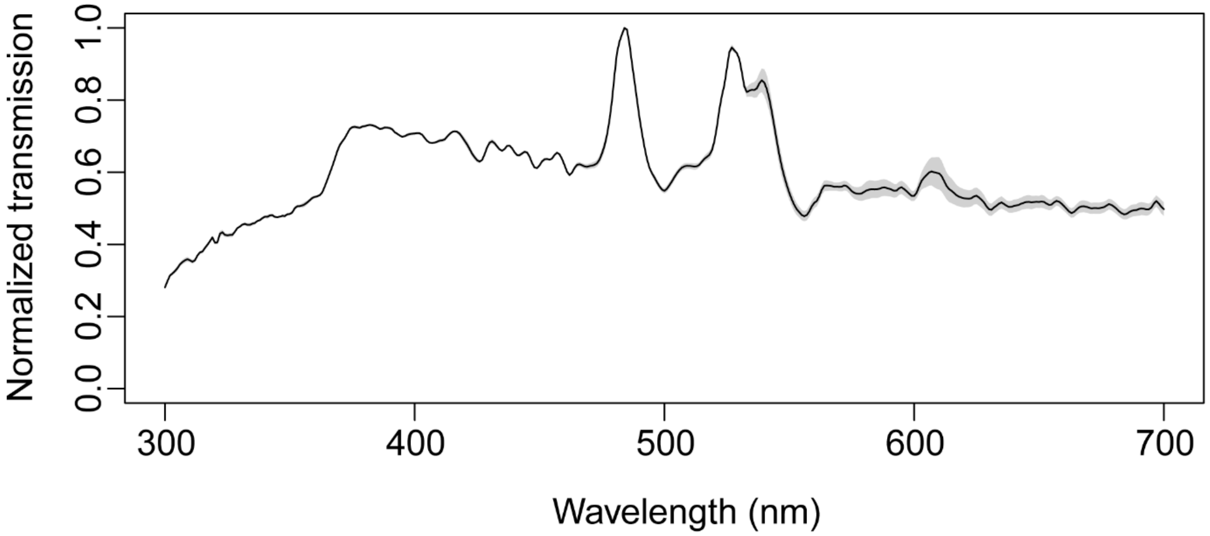


**Supplementary Figure 8.** Normalized transmission spectra measured through the clear plastic of the juvenile containers used in the behavior experiments. Plotted is the average transmission spectra (n = 3) with standard error shaded in grey.
