## Supplementary material for "Socially regulated developmental plasticity in the color pattern of an anemonefish": Figure S9

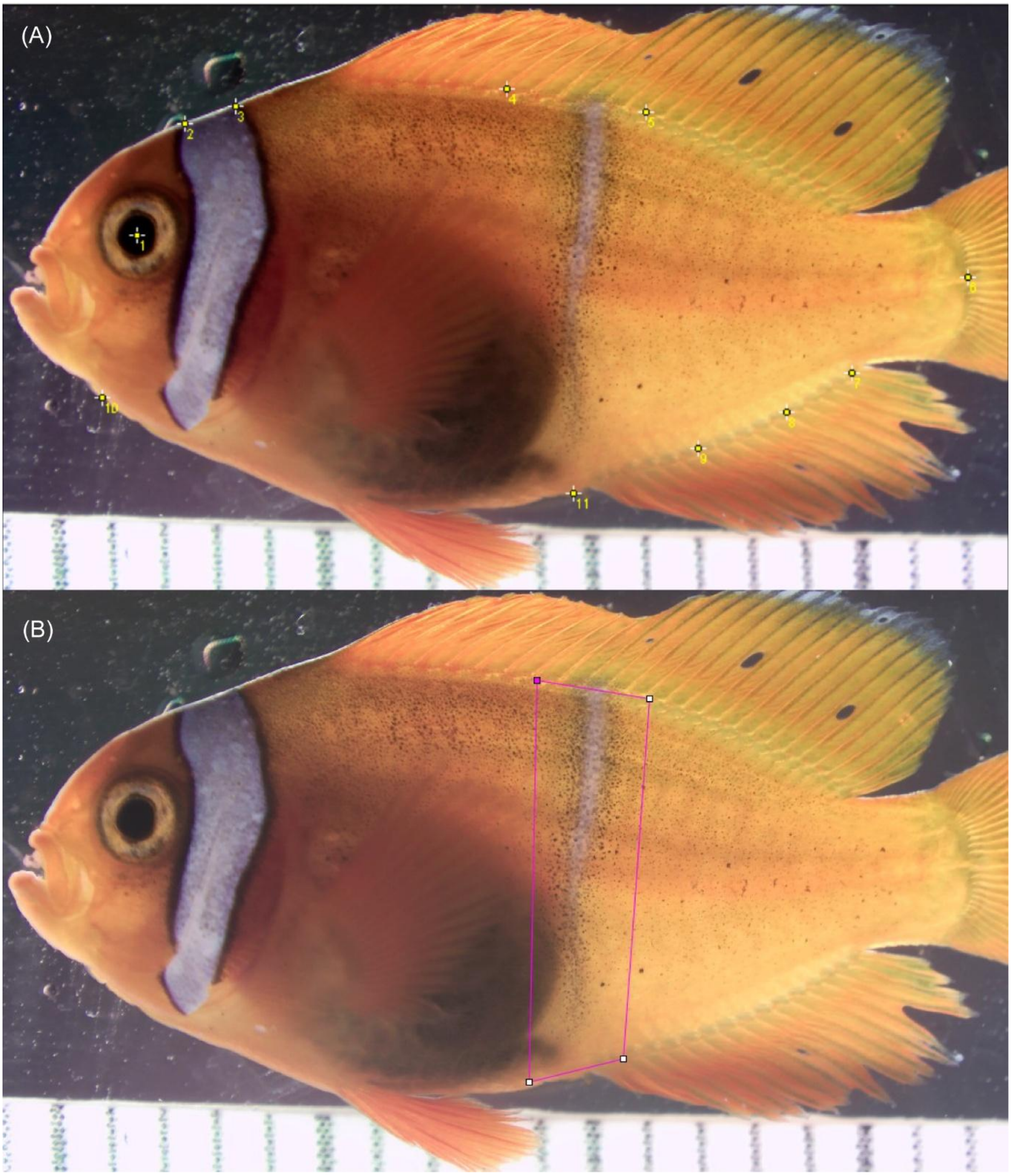


**Supplementary Figure 8. (A)** Landmark locations used for the alignment of fish images in colour pattern area analysis. **(B)** Vector locations for the mask used for isolating the body bar region were dorsally at the 8$^{th}$ dorsal spine and 4$^{th}$ dorsal ray, and  in-line ventrally with the 2$^{nd}$ anal fin spine and posterior-edge of the pelvic fin.
