## Supplementary material for "Socially regulated developmental plasticity in the color pattern of an anemonefish": Table S1

### Supplementary Table 1. Ordbeta reg summary output for white skin proportion in development experiment

| | *Estimate* | *Std. Error* | *z value* | *Pr(>\|z\|)* |
|---|---|---|---|---|
| *Intercept* | -3.66 | 0.34 | -10.80 | <2e-16 |
| *treatment_anemone* | 1.55 | 0.36 | 4.36 | 1.30e-05 |
| *treatment_fakeanemone* | 2.31 | 0.31 | 7.38 | 1.58e-13 |
| *treatment_empty* | 2.24 | 0.32 | 7.05 | 1.75e-12 |
| *sampled_age* | -0.76 | 0.81 | -0.95 | 0.34 |
| *sl_mm* | -0.56 | 0.14 | -3.92 | 9.02e-05 |
| *treatmentB:sampled_age* | -1.36 | 0.95 | -1.44 | 0.15 |
| *treatmentC:sampled_age* | 0.93 | 0.79 | 1.17 | 0.24 |
| *treatmentD:sampled_age* | 0.98 | 0.80 | 1.23 | 0.22 |
