## Supplementary material for "Socially regulated developmental plasticity in the color pattern of an anemonefish": Table S2

**Supplementary Table 2. Age grouped comparisons with adjusted coefficients for development experiment**

| *treatment* | *response* | *SE* | *LCL* | *UCL* |
|---|---|---|---|---|
| *A:38dph* | 0.03 | 0.008 | 0.013 | 0.05 |
| *B:38dph* | 0.11 | 0.02 | 0.08 | 0.15 |
| *C:38dph* | 0.21 | 0.04 | 0.15 | 0.28 |
| *D:38dph* | 0.20 | 0.03 | 0.14 | 0.26 |
| *A:62dph* | 0.01 | 0.009 | 0.003 | 0.05 |
| *B:62dph* | 0.01 | 0.01 | 0.003 | 0.06 |
| *C:62dph* | 0.24 | 0.03 | 0.18 | 0.30 |
| *D:62dph* | 0.23 | 0.03 | 0.18 | 0.30 |

‘*LCL*’ = lower confidence limit; ‘*UCL*’ = upper confidence limit.
