## Supplementary material for "Socially regulated developmental plasticity in the color pattern of an anemonefish": Table S3

**Supplementary Table 3.** GLMM output summary for Behavior Experiment 1.

| | *Estimate* | *Std. error* | *z value* | *Pr(>\|z\|)* |
|---|---|---|---|---|
| *(intercept)* | 1.06 | 0.14 | 7.54 | 4.83e-14*** |
| *bar_no* | -2.04 | 0.19 | -10.78 | <2e-16*** |
| *delta_SL* | 0.17 | 0.11 | 1.60 | 0.11 |

trial_ID: variance = 0.095, Std. Dev. = 0.31; tank_ID: variance = 1.16e-10, Std. Dev. = 1.08e-05.
