## Supplementary material for "Socially regulated developmental plasticity in the color pattern of an anemonefish": Table S4

**Supplementary Table 4.** GLMM output summary for Behavior Experiment 2.

| | *Estimate* | *Std. error* | *z value* | *Pr(>\|z\|)* |
|---|---|---|---|---|
| *(intercept)* | -1.48 | 0.32 | -4.63 | 3.69e-06*** |
| *delta_SL* | 0.33 | 0.17 | 1.91 | 0.06 |
| *bar_no* | 2.56 | 0.31 | 8.39 | <2e-16*** |

trial_ID: variance = 5.21e-10, Std. Dev. = 2.28e-05; tank_ID: variance = 0.18, Std. Dev. = 0.42.
